# The draft genome of the invasive walking stick, *Medauroidea extradendata*, reveals extensive lineage-specific gene family expansions of cell wall degrading enzymes in Phasmatodea

**DOI:** 10.1101/285817

**Authors:** Philipp Brand, Wei Lin, Brian R. Johnson

## Abstract

Plant cell wall components are the most abundant macromolecules on Earth. The study of the breakdown of these molecules is thus a central question in biology. Surprisingly, plant cell wall breakdown by herbivores is relatively poorly understood, as nearly all early work focused on the mechanisms used by symbiotic microbes to breakdown plant cell walls in insects such as termites. Recently, however, it has been shown that many organisms make endogenous cellulases. Insects, and other arthropods, in particular have been shown to express a variety of plant cell wall degrading enzymes in many gene families with the ability to break down all the major components of the plant cell wall. Here we report the genome of a walking stick, *Medauroidea extradentata*, an obligate herbivore that makes uses of endogenously produced plant cell wall degrading enzymes. We present a draft of the 3.3Gbp genome along with an official gene set that contains a diversity of plant cell wall degrading enzymes. We show that at least one of the major families of plant cell wall degrading enzymes, the pectinases, have undergone a striking lineage-specific gene family expansion in the Phasmatodea. This genome will be a useful resource for comparative evolutionary studies with herbivores in many other clades and will help elucidate the mechanisms by which metazoans breakdown plant cell wall components.

**Data availability:** The *Medauroidea extradentata* genome assembly, Med v1.0, is available for download via NCBI (Bioproject: PRJNA369247). The genome, annotation files, and official gene set Mext_OGS_v1.0 are also available at the i5k NAL workspace (https://i5k.nal.usda.gov/medauroidea-extradentata) and at github (https://github.com/pbrec/medauroidea_genome_resources). The genomic raw reads are available via NCBI SRA: SRR6383867 and the raw transcriptomic reads are available at NCBI SRA: SRR6383868, SRR6383869.

## Introduction

The components of the plant cell wall are the most abundant macromolecules on earth and the study of their breakdown by herbivores and decomposers is thus of central importance to biology (Beguin and Aubert 1994; Keegstra 2010). Plant cell walls (PCWs) contain lignocellulosic compounds that are difficult to degrade, such as xylan, cellulose, hemicellulose, pectin, and lignin (Cosgrove 2005). Degradation of PCWs requires the ability to physically degrade the tough material, then biochemically breakdown some or all of its components (Calderon-Cortes et al 2012). Organisms across the tree of life employ a diverse set of strategies to accomplish this with some able to utilize all PCW components and others only a subset. Central to all approaches are plant cell wall degrading enzymes (PCWDEs) falling into several gene families (Beguin and Aubert 1994; Lo et al 2003; Watanabe and Tokuda 2010). In addition to being of interest to those studying ecology and physiology, PCWDE’s are also of interest to those in the biofuel industry, as the efficient breakdown of cellulose to simple sugars is central to the utility of biofuels (Pauly and Keegstra 2010).

Invertebrates, chiefly insects, are major herbivores and decomposers in many ecosystems and effectively use lignocellulosic materials for energy. Early work on termites, the major group of strictly wood feeding insects, suggested that PCWDEs produced by bacterial symbionts are required for insect breakdown of PCWs (Martin 1991; Breznak and Brune, 1994). This was supported both by studies of microbes in termites, but also by work on model systems, flies and butterflies, that showed a lack of symbionts and a lack of PCW breakdown ability (Slaytor 1992; reviewed in Watanabe and Tokuda 2010). Recent work, however, has shown that endogenously produced PCWDEs are more widespread and important in insects than previously thought (Watanabe et al 1998; Lo et al 2003; Nakashima et al 2002; Shelomi et al 2014a,b; Bai et al 2016; Wu et al 2016). First, a closer examination of termites showed that they also produce endogenous PCWDEs, and second, studies of other insects showed widespread production of endogenously produced PCWDEs. A current obstacle to understanding the diversity of PCWDE’s in insects is sampling bias in the sequencing of genomes, towards holometabolous insects. It is likely many holometabolous insects lack the diversity of PCWDE’s present in some clades of hemimetabolous insects (reviewed in Watanabe and Tokuda 2010). This prediction is based on the discovery that so far only Coleoptera and Hymenoptera in the Holometabola have been found to have PCWDEs (and only Coleoptera in large numbers), while most hemimetabolous insects sequenced thus far have them, including several clades with extensive repertoires (reviewed in Watanabe and Tokuda 2010).

Phasmids, walking sticks, are large long-lived insects that feed exclusively on leaves. Previous work using transcriptomics has shown that phasmids express a diversity of PCWDEs, including cellulases, hemicellulases, and pectinases (Shelomi et al 2014a,b, 2016; Wu et al 2016). This work also suggested that gene duplications in the cellulases have led to enzymes with the capacity to break down multiple components of the PCW (Kirsh et al 2014; Shelomi et al 2016). Of further interest, pectinases found in more derived phasmid transcriptomes is more similar to bacterial pectinases than to those known from eukaryotes, suggesting horizontal gene transfer (Shelomi et al 2016b). Such work highlights the utility of phasmids as models for the study of PCW breakdown evolution.

Here we present a draft genome for *Medauroidea extradentata*, a common invasive walking stick found in many parts of the world. The ease of culturing these insects in the lab, and their widespread distribution, makes them a suitable potential model system for laboratory studies of PCWDEs. We used the DISCOVAR approach coupled with RNA-Seq based scaffolding to produce the draft genome. Annotation of the genome, assisted by several RNA-Seq datasets, produced a high-quality gene set comparable to those of other sequenced invertebrates with large genome size. Analyses of the pectinase gene family, in comparison to those pectinases in other hemimetabolous insects, supports a single horizontal gene transfer event of pectinase genes from bacteria to Phasmatodea. In addition, we identify more extensive than previously thought lineage-specific expansions of this gene family following the horizontal transfer event.

## Materials and Methods

### Genome sequencing and assembly

DNA was extracted from a single female wild caught *Medauroidea extradentata* adult, captured near Sacramento CA, with a Qiagen DNeasy kit using manufacturer’s instructions. The digestive tract was first removed from the insect to minimize contamination from food items and microbes. DNA was tested for purity with the nanodrop 1000 and for concentration with the Qubit 3.0. A single sequencing library was then made with the Truseq DNA PCR Free library preparation kit according to the manufacturer’s instructions. The library was quality tested with the Bioanalyzer 2000 and 250 base pair paired end sequencing was conducted on the Illumina Hiseq 2500. A total of 355,738,482 reads were produced. Assembly of the resulting reads was performed with DISCOVAR (version 1) using default parameters (Weisenfeld et al 2014).

Because the sample was wild caught, it can be expected to be more heterozygous than is typical for genome studies which often use inbred lab strains. Accordingly, to reduce assembly errors due to high heterozygosity, Redundans (version 1), with default parameters (contigs with greater than 85% similarity to other longer contigs removed), was used to reduce the number of duplicate contigs from the initial DISCOVAR assembly (Pryszcz and Gabaldon 2016). Because the resulting assembly was still fragmented, a final scaffolding step was performed with Agouti (version 1), an RNA-Seq based scaffolder, using default parameters (Zhang et al 2016). This assembly was labeled “Med v1.0” and was used for all subsequent analyses. Earlier assemblies can be produced from the raw reads, available at NCBI, or are available upon request. The quality of the assembly was assessed using busco v2 (Simao et al 2015). Busco was run using the arthropoda_odb9 database in genome mode.

### Genome size estimation

Genome size was estimated from the sequencing reads based on the k-mer frequency spectrum. A k-mer library based on all sequence reads was prepared using Jellyfish with a k value of 25. The resulting k-mer frequency spectrum was then used to estimate genome size on the basis of the consecutive length of all reads divided by the sequencing depth as previously described (Brand et al. 2017).

### Genome annotation

Several libraries were constructed and sequenced to facilitate genome annotation. In short, RNA was extracted from freshly dissected tissue with Trizol and quality controlled with the Bioanalzyer 2100 to ensure no degradation. Quantification of RNA was done with the Qubit 3.0. All libraries were 150 bp PE and were constructed with the NEBNext® Ultra™ RNA Library Prep Kit for Illumina using the manufacturer’s instructions. All samples were from female insects. RNA was extracted from: whole bodies of juvenile insects (5 pooled insects), the reproductive tract of adults (5 pooled insects), and 3 pools of 5 insects for the Malpighian tubules of adult insects. In addition, a previous study provided 3 libraries of the anterior midgut and 3 libraries of the posterior midgut (Shelomi et al 2014a). Separate libraries were produced for juveniles and adults, but the resulting reads were pooled for transcriptome assembly using Bridger (r2014-12-01) with default parameter settings (Chang et al 2015). In total ∼170 million 150 PE reads were produced for juvenile whole body libraries, ∼50.5 million 150 PE for adult reproductive tracts, and ∼152 million 150 PE reads total for the Malpighian tubules.

Maker was used for annotation with commonly used recommended settings (Cantarel et al 2008). In short, Augustus was used for *ab initio* gene prediction (Stanke et al 2004) using the training from aphid (nearest insect from available options), blastx was used for protein homology searches, and tblastn was used to align cDNAs from the transcriptome to the genome. Repeat masker was used to mask repetitive DNA during annotation. We provide a high-confidence subset of all gene annotations based on gene expression quantification and homology to the stick insect *Timema cristinae*. We identified reciprocal best blast hits (BBH) between our gene models and *T. cristinae* (Soria-Carrasco et al. 2014) using blastp with an evalue-cutoff of 10E-12. In addition, we used Kallisto (Bray et al. 2016) to infer expression levels of all gene models based on our RNA-Seq libraries. All genes with a BBH to *T. cristinae* and/or a TPM (transcripts per million) estimate ≥ 1 were included in the high-confidence gene set, representing the *M. extradentata* official gene set (Mext_OGS_v1.0).

### Repetitive element annotation

#### Tandem repeats

Micro- and mini-satellites (1-6 bp and 7-1000 bp motif length, respectively) were annotated in all scaffolds ≥1000bp using Phobos 3.3.12 (Mayer 2010). Therefore, one independent run for each class of tandem repeats was performed with Phobos parameter settings following (Leese et al. 2012: gap score and mismatch score set to −4 and a minimum repeat score of 12).

#### TEs

In order to annotate TEs, RepeatModeler was used for de novo repeat element annotation and classification followed by RepeatMasker to detect the total fraction of repetitive elements present in the genome assembly (Smit et al. 2016). RepeatModeler v1.0.8 was run with default settings using the NCBI blast algorithm (Altschul et al. 1990) for repeat detection. The resulting de novo TE annotations were used as a database for Repeatmasker v4.0.5 with Crossmatch in the sensitive mode. Low complexity regions were excluded from the analysis.

Plant cell wall degrading enzyme annotation and phylogenetic analysis A combination of tblastn and exonerate (Altschul et al. 1990; Slater and Birney 2005) was used to manually annotate genes of the pectinase [polygalacturonase] and cellulase [endo-beta-1,4-glucanase] gene families. We used the semi-automated pipeline described in Brand and Ramirez (2017). Briefly, genes known from bacteria and eukaryotes including fungi, plants, and insects were used as query to identify scaffolds with significant tblastn hits (e-value <10E-6). Subsequently, we used exonerate to identify potential intron-exon boundaries of genes on the respective scaffolds. Resulting gene models with a minimum length of 150 amino acids were included in the gene families. In addition to *M. extradentata*, we annotated three phasmids *Dryococelus australis* (Mikheyev et al. 2017), *Clitarchus hookery* (Wu et al. 2017) and *Timema cristinae* (Riesch et al. 2017), as well as the German cockroack *Blatella germanica* (Harrison et al. 2018) and the termite *Zootermopsis nevadensis* (Terrapon et al. 2014).

To identify the putative evolutionary origin of the annotated pectinase gene sequences, we used available annotations to check for eukaryote gene models in the 20kb flanking regions of each gene upstream and downstream in the respective genomes. If no gene models containing multiple exons were located within the flanking regions, we used blastn against the NCBI nuccore database to identify the origin of the gene. Using an evalue threshold of 10E-6, genes were either identified as of insect, bacterial, or unknown origin.

In order to understand the evolutionary history of the two PCWDE gene families, we next inferred the gene family phylogenies. Therefore, the protein sequences of the identified genes of all six hemimetabolous insects and genes from outgroups covering main bacterial and all major eukaryote lineages (GIs from Shelomi et al. 2014a) were used to produce an alignment using mafft (Katoh et al. 2002) applying the L-INS-I algorithm with the --maxiterate option set to 1,000. The alignments were manually examined for conserved functional sites (Shelomi et al. 2014a) and used for maximum likelihood gene tree inference with RaXML (Stamatakis et al. 2005) using the JTT + gamma substitution model.

#### Data availability

The *Medauroidea extradentata* genome assembly, Med v1.0, is available for download via NCBI (Bioproject: PRJNA369247). The genome, annotation files, and official gene set Mext_OGS_v1.0 are also available at the i5k NAL workspace (https://i5k.nal.usda.gov/medauroidea-extradentata) and at github (https://github.com/pbrec/medauroidea_genome_resources). The genomic raw reads are available via NCBI SRA: SRR6383867 and the raw transcriptomic reads are available at NCBI SRA: SRR6383868, SRR6383869.

## Results and Discussion

### Basic assembly and annotation

Genome size was estimated to be 3.3Gbp based on the kmer analysis (Table 1). The *Medauroidea extradentata* genome assembly has 135,692 scaffolds with an N50 score of 43,047 (Table 1). The final genome assembly (post redundans and post agouti) is 2.6Gbp which is 78.8% percent of the estimated size based on kmer counts. Coverage was found to be approximately 54-fold based on the total amount of DNA produced and the estimated genome size. Genomic GC content was 37%.

**Table 1.**
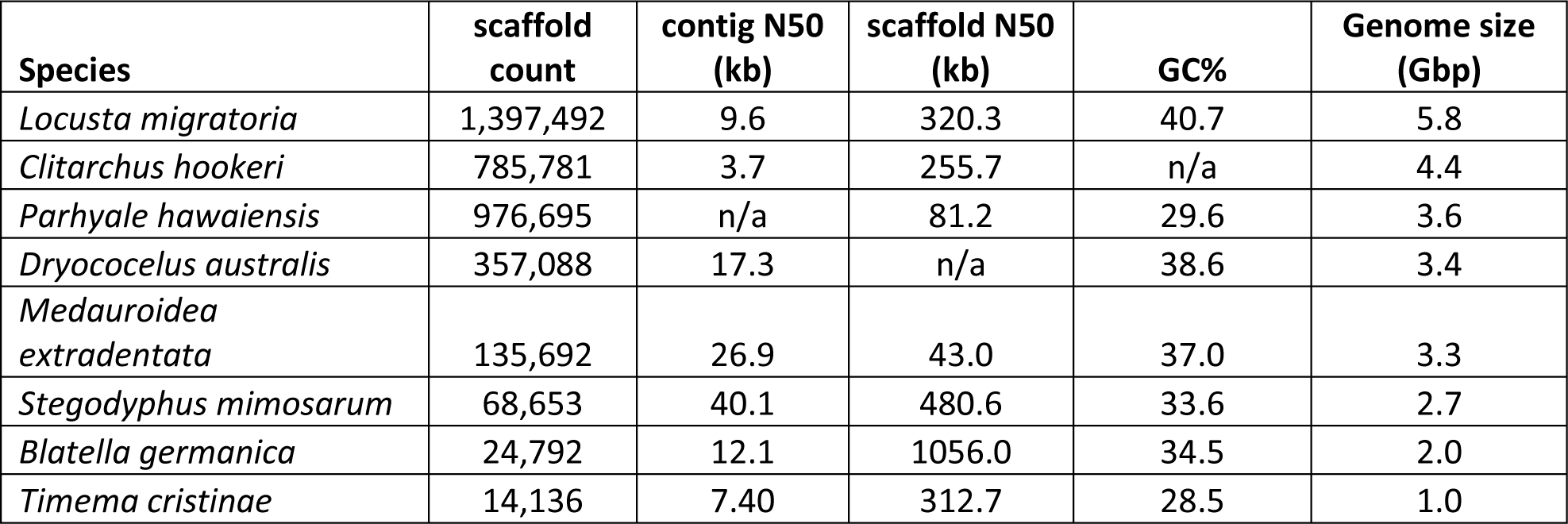
Basic assembly statistics for several recently sequenced large genome arthropods (Wang et al 2014; Soria-Carrascoet al 2014; Sanggaard et al 2014; Kao et al 2016; Harrison et al 2017; Wu et al 2017; McGrath et al 2017).

| **Species** | **scaffold count** | **contig N50 (kb)** | **scaffold N50 (kb)** | **GC%** | **Genome size (Gbp)** |
| --- | --- | --- | --- | --- | --- |
| *Locusta migratoria* | 1,397,492 | 9.6 | 320.3 | 40.7 | 5.8 |
| *Clitarchus hookeri* | 785,781 | 3.7 | 255.7 | n/a | 4.4 |
| *Parhyale hawaiiensis* | 976,695 | n/a | 81.2 | 29.6 | 3.6 |
| *Dryococelus australis* | 357,088 | 17.3 | n/a | 38.6 | 3.4 |
| *Medauroidea extradentata* | 135,692 | 26.9 | 43.0 | 37.0 | 3.3 |
| *Stegodyphus mimosarum* | 68,653 | 40.1 | 480.6 | 33.6 | 2.7 |
| *Blatella germanica* | 24,792 | 12.1 | 1056.0 | 34.5 | 2.0 |
| *Timema cristinae* | 14,136 | 7.40 | 312.7 | 28.5 | 1.0 |

The Busco analysis showed a level of completeness comparable to that for other large arthropod genomes (Table 2). 78.8% of genes in the Arthropod DB were complete, 17.4% were present but fragmented and only 3.8% were missing. For comparative purposes, two large arthropod genomes recently published (Parhyale, a crustacean, and Locust) have values of 78.5% and 41.4% for completeness, 10.4% and 31.5% for fragmented, and 11.1% and 27.1% for missing (Wang et al 2014; Kao et al 2016). Essentially, these 3 large genomes are 10 times the size of most holometabolous insect genomes (which are about 300MB) and have very high levels of repetitive DNA (Kidwell 2002). It is thus not surprising that they are less complete at the first draft stage, though future work should be conducted to improve these assemblies.

**Table 2.** Busco analysis comparison for other large genome size arthropods. Complete refers to genes within a core list of one to one orthologs across the arthropods (arthopoda_db) that are complete in the present assembly. Fragmented and missing likewise refer to highly conserved genes from arthopoda_db that are either present, but incomplete (fragments), or missing from the present assembly.

| **Species** | **Complete** | **Fragmented** | **Missing** |
| --- | --- | --- | --- |
| *Locusta migratoria* | 41.4 | 31.5 | 27.1 |
| *Parhyale hawaiiensis* | 78.5 | 10.4 | 11.1 |
| *Medauroidea extradentata* | 78.8 | 17.4 | 3.8 |
| *Stegodyphus mimosarum* | 92.1 | 2.7 | 5.2 |
| *Blatella germanica* | 98.8 | 0.7 | 0.5 |

**Table 3.** Results of transposable element repeat class analysis.

| **Repeat Element Family** | **Number Unique Elements** | **Total Number Elements in Assembly** | **Cumulative Length (bp)** | **Percent of Genome Assembly<sup>a</sup>** |
| --- | --- | --- | --- | --- |
| **Class I - retrotransposons** | 141 | 547,153 | 180,544,125 | 6.98 |
| **Class II - DNA transposons** | 171 | 354,700 | 124,099,554 | 4.80 |
| **Total classified transposons** | 312 | 901,853 | 304,643,679 | 11.79 |
| **Unclassified** | 1,097 | 3,323,694 | 969,506,662 | 37.51 |
| **Total repeat families** | 1,409 | 4,225,547 | 1,274,150,341 | 49.29 |

Our annotation efforts resulted in a total of 103,773 preliminary gene models of which 35,742 were homologous to *T. cristinae* and/or expressed based on RNA-Sequencing and thus constitute the official gene set (OGS version 1.0).

#### Repetitive elements

##### Tandem Repeats

A total of 673,636 microsatellite loci with a consecutive length of 23,936,685 bp were detected. Minisatellites with motif lengths from 7 bp to 1000 bp were less numerous in the genome with 257,457 loci but had a higher accumulative length (44,403,552 bp). Accordingly, tandem repeats represent 2.6% of the assembly, suggesting that they contribute a small proportion to the overall genome size.

##### TEs

The RepeatModeler analysis revealed a total of 1409 repeat element families in the assembly of which 312 (22.1%) belonged to known TE families including 171 DNA transposons and 141 retroelements. The remaining 1097 (77.9%) repeat element families could not be classified into known TE families. All 1409 detected repeat element families were used as database for the RepeatMasker analysis. This way, a total of 4,225,547 elements were annotated in the assembly of which 901,853 (21.3%) were derived from the 312 classified TE families. The remaining 3,323,694 (78.7%) elements belonged to the unclassified repeat element families. In total, all annotated repeat elements had a cumulative length of 1,274,150,341 bp corresponding to 49.29% of the total genome assembly length. The majority of repeat elements were derived from unclassified families corresponding to 37.51% of the total assembly length.

Given the large genome size, the detected high fraction of the genome associated with repetitive element families is not surprising. Large genome sizes in insects and most other organisms are generally associated with elevated TE activity and content (Kidwell 2002). Although this correlation is ubiquitous in nature, most repetitive elements associated with this form of ‘genome obesity’ are not very well characterized, due to the fast evolving nature of TEs, which leads to large underestimates of genomic TE content in non-model lineages (Chalopin et al. 2015; Platt et al. 2016). This likely explains the large fraction of unclassified repetitive element families detected in the present genome assembly. In total, our analysis suggests that a large proportion of the *M. extradentata* genome is repetitive. This result is similar to other insects with comparable genome sizes (Wang et al. 2014, Brand et al. 2017).

##### Plant cell wall degrading enzymes and the evolution of pectinases and cellulases

*M. extradentata* was chosen for genome sequencing due to its potential use as a model system for studies of the physiology of herbivory, particularly plant cell wall breakdown. PCWDEs were present in large numbers in the *M. extradentata* genome (5 cellulase gene models, 87 pectinase gene models, 3 beta-1,3-glucanase gene models, and 33 cellobiase gene models). A detailed analysis of cellulases and pectinases across six hemimetabolous insect herbivores revealed large variation in the size of the pectinase family, and less but still significant variation in the size of the cellulase gene family (Figure 1). While cellulases were present in all species analyzed, we only identified pectinases in the Phasmatidae (*M. extradentata, C. hookeri, D. australis*) and the cockroach *B. germanica.*

**Figure 1.**
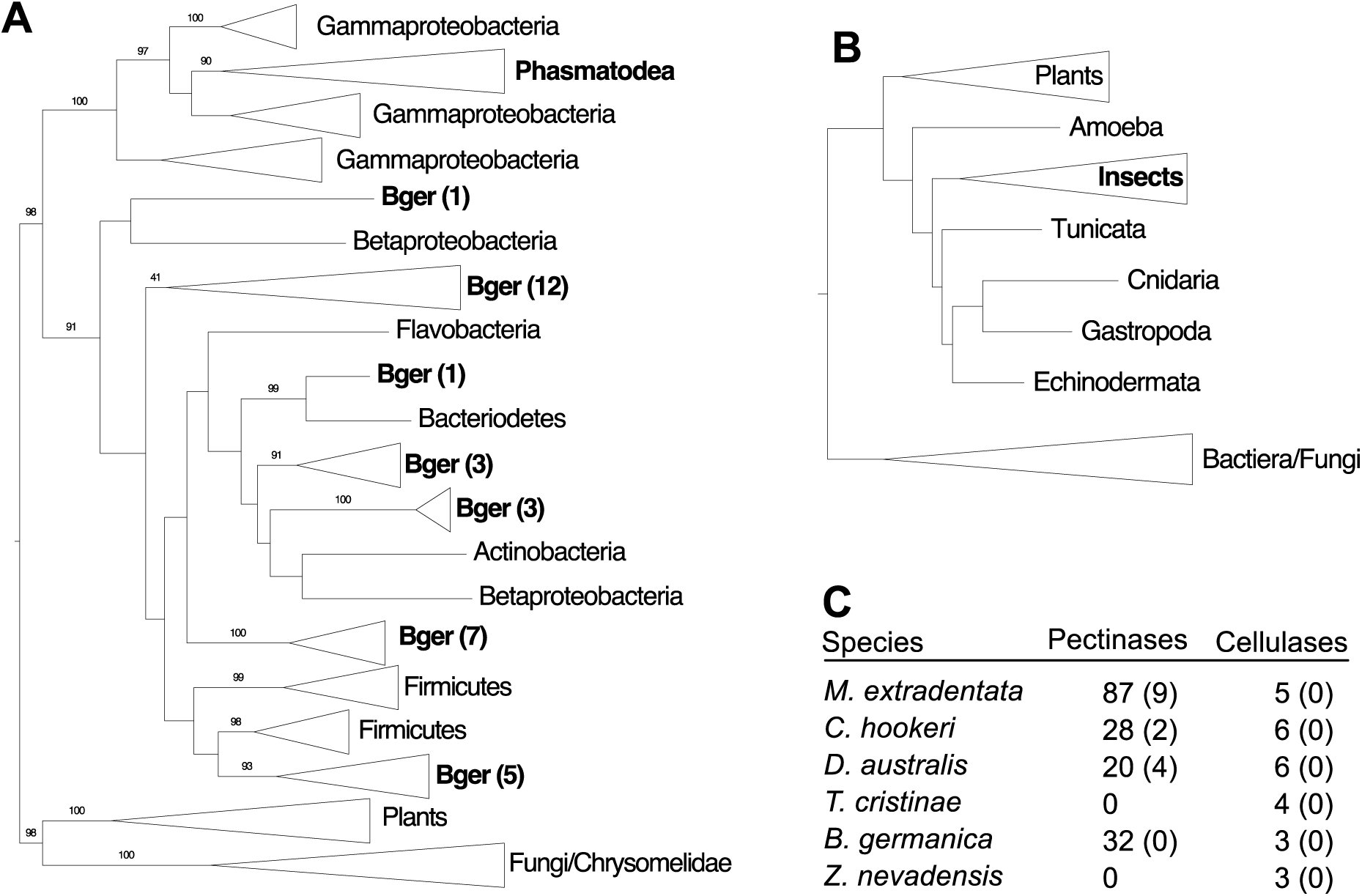
Cell wall degrading enzyme gene family dynamics. **A)** The pectinase genes identified in the three Phasmatodea species all clustered within the gammaproteobacteria, while the pectinases identified in the *B. germanica* genome (Bger) were located throughout the bacteria. Numbers in brackets indicate the number of *B. germanica* genes in collapsed clades. **B)** All cellulase genes identified clustered in a single insect clade. **C)** Table including pectinase and cellulase genes identified in the six hemimetabolous species. Numbers in brackets represent the number of pseudogenes. Bootstrap support ≥ 90 are marked on respective branches.

Gene family specific phylogenies showed that all identified pectinases were more closely related to bacterial than eukaryotic pectinases (Figure 1A). Nevertheless, most pectinases in the genomes of the Phasmatidae species were located near eukaryotic genes, suggesting that they were inserted in the insect genome and not due to bacterial contamination. While the pectinases in the cockroach were similarly located in large scaffolds with eukaryote gene predictions, the 20kb flanking regions never contained eukaryotic genes and were more similar to bacterial than insect genomic sequences. Accordingly, we were not able to unequivocally identify if the genes were located on the insect genome or part of bacterial contamination leading to genome assembly artifacts.

Interestingly, all phasmatida pectinases clustered as a monophyletic group within the gammaproteobacteria clade. We identified seven 1:1 orthologous pectinase genes in *C. hookeri* and *D. australis*, as well as 7 duplications or larger expansions specific to *C. hookeri* (Supplemental Figure 1). *M. extradentata* on the other hand had only 4 pectinases with simple 1:1 or 1:1:1 orthology to pectinases of the other two species. Most pectinases detected in the *M. extradentata* genome were part of large lineage-specific gene family expansions. These results confirm that a single horizontal gene transfer from gammaproteobacteria preceding the split of the Phasmatidae is the most likely mechanism for the origin of pectinase genes in the genome of this insect lineage (Shelomi et al. 2016b), and that pectinases evolved through a birth-death mechanism common for multi-gene families (Nei and Rooney 2005) after the horizontal gene transfer event.

Similar to the Phasmatidae, the pectinases detected in the cockroach genome were more closely related to bacteria than eukoryotes, however, they clustered with multiple different bacterial lineages (Supplementary Figure 1). This suggests different bacterial origins of the pectinases associated with the two lineages of hemimetabolous insects. In contrast to the Phasmatidae, these findings do not support a single horizontal gene transfer event from bacteria to cockroaches, but rather indicate that the identified pectinases are indeed of bacterial origin. It is likely that the identified pectinases in the genome assembly represent assembly artifacts due to bacterial contamination. Accordingly, the origin of the pectinases identified in the cockroach genome needs to be verified in future hemiptera-specific analyses.

In comparison to the pectinases, the cellulase gene family was more similar between species. All cellulases clustered within insects in lineage-specific clades (Figure 1B; Supplemental Figure 2).

## Conclusions

The large 3.3Gbp *Medauroidea extradentata* genome presented here will facilitate the further exploration of the evolution of PCW breakdown in phasmids, a complex process involving numerous gene duplications and horizontal gene transfer. The large gene family for pectinases, in particular, which varies strongly in size across the Phasmatodea and other insect orders, will be a promising candidate for future work. Further, hemimetabolous insects, and phasmids in particular, are still poorly represented in genome studies; this work therefore contributes to a more balanced representation of available genomes for evolutionary studies. Finally, this work will also facilitate studies of repetitive element evolution, as there is slowly building up a sufficiently large number of large arthropod genomes for comparative analysis in this context.

## Table and Figure Legends

**Supplemental Figure 1 Phylogenetic tree of the pectinase gene family.** All phasmatodea pectinases cluster within the gammaproteobacteria and form a highly supported monophyletic clade, supporting a single horizontal gene transfer event in the ancestor of the three insect lineages analyzed. Pectinases detected in the *B. germanica* genome cluster within bacteria as well, but do not form a monophyletic clade. It is likely that the pectinases identified are due to bacterial contamination of the genome assembly (see main text). Known chrysomelid beetle pectinases cluster within fungi, representing independent horizontal gene transfer events of pectinases from fungi to insects (Pauchet et al. 2010) Orange: *M. extradentata*, Blue: *C. hookeri*, Purple: *D. australis*, Red: *B. germanica.* Bootstrap support of 100 replicates is indicated for each branch. Gene models of the four newly annotated species with insect or bacterial genes in the 20kb flanking regions are indicated.

**Supplemental Figure 2 Phylogenetic tree of the cellulase gene family.** All identified cellulase genes cluster within other known bacterial cellulase genes. Orange: *M. extradentata*, Blue: *C. hookeri*, Purple: *D. australis*, Red: *B. germanica*, Brown: *Z. nevadensis*, Pink: *T. cristinae*. Bootstrap support of 100 replicates is indicated for each branch.

